# Pyocin efficacy in a murine model of *Pseudomonas aeruginosa* sepsis

**DOI:** 10.1101/2020.03.27.011908

**Authors:** Anne Six, Khedidja Mosbahi, Madhuri Barge, Colin Kleanthous, Thomas Evans, Daniel Walker

## Abstract

**Background:** Bloodstream infections with antibiotic resistant *Pseudomonas aeruginosa* are common and increasingly difficult to treat. Pyocins are naturally occurring protein antibiotics produced by *P. aeruginosa* that have potential for human use.

**Objectives:** To determine if pyocin treatment is effective in a murine model of sepsis with *P. aeruginosa*.

**Methods:** Recombinant pyocins S5 and AP41 were purified tested for efficacy in a *Galleria mellonella* infection model and a murine model of *P. aeruginosa* sepsis.

**Results:** Both pyocins produced no adverse effects when injected alone into mice and showed good in vitro antipseudomonal activity. In an invertebrate model of sepsis using *Galleria mellonella*, both pyocins significantly prolonged survival. Following injection into mice, both showed extensive distribution into different organs. When administered 5 hours after infection, both pyocins reduced mortality, with pyocin S5 being more effective than AP41.

**Conclusions:** Pyocins S5 and AP41 show in vivo biological activity and can improve survival in a murine model of *P. aeruginosa* infection. They hold promise as novel antimicrobial agents for treatment of multi-drug resistant infections with this microbe.

## Introduction

*Pseudomonas aeruginosa* is a leading cause of severe hospital-acquired infections such as ventilator-associated pneumonia, burn wound infections and nosocomial bloodstream infections (BSI). *P. aeruginosa* accounts for 4% of all hospital-acquired BSI cases and has a very high (>30%) mortality rate as well as high healthcare costs^1–7^. In addition, because of its limited susceptibility to antibiotics and the frequent emergence of resistance during therapy ^8,9^. *P. aeruginosa* infections are difficult to treat. In common with many bacterial pathogens, there is an increasing prevalence of multidrug-resistant strains, leading to the classification of *P. aeruginosa* as critical on the WHO list of antibiotic-resistant priority pathogens^10^. In Europe, the 2017 report of the ECDC showed that 30.8% of *P. aeruginosa* were resistant to at least one of the antimicrobial groups under surveillance^11^.

A potential alternative therapeutic strategy to treat multidrug-resistant is the use of the highly potent and narrow spectrum bacteriocins as protein antibiotics^12^. Bacteria produce these very diverse and widespread proteins during intraspecies competition, where they kill only bacteria closely related to the producing strain^13,14^ Bacteriocins from *P. aeruginosa* can be classified into different categories, with the most abundant S-type pyocins closely resembling colicins from *Escherichia coli*^14,15^. These multi-domain proteins share similar structural organization (including a receptor binding domain, a translocation domain and a cytotoxic domain), and are able to efficiently cross the Gram-negative outer membrane through parasitization of nutrient uptake pathways^16^. Indeed, the uptake pathways for several pyocins from *P. aeruginosa* have been identified and these typically involve outer membrane transporters involved in the uptake of iron siderophores^17–21^. The cytotoxic activity of pyocins generally takes the form of a nuclease activity targeting DNA (such as pyocin AP41), tRNA or rRNA, a pore-forming activity targeting the cytoplasmic membrane (pyocin S5) or an enzymatic activity targeting peptidoglycan synthesis^22^.

The potency and specificity pyocins makes them potential candidates for the treatment of *P. aeruginosa* infection. We previously demonstrated pyocin efficacy for the treatment of acute pneumonia in a murine model^23^. In this work, we show that pyocin S5 and pyocin AP41 delivered directly to the blood retain their bactericidal activity in vivo and display efficacy against a clinical strain of *Pseudomonas aeruginosa* in a murine sepsis model of infection. Furthermore, we show using a *Galleria mellonella* systemic infection model, that pyocin S5 is able to treat a high-level antibiotic-resistant infection, highlighting their potential as therapeutics in a world of increasing antimicrobial resistance.

## Materials and methods

### Bacterial strains, genetic constructions and growth conditions

The plasmid pETPyoAP41, which encodes the genes for pyocin AP41 and the C-terminally His_6_-tagged immunity protein, ImAP41, was used for expression of the pyocin AP41-ImAP41 complex as previously described^23^. pPyoS5 was used for the expression of pyocin S5 (with no affinity tag). This plasmid was constructed by ATUM and encodes the pyocin S5 gene, optimised for expression in *E. coli* in the vector pJ404. *Pseudomonas aeruginosa* P7 strain is a clinical mucoid strain isolated from a CF patient^23^. *E. coli* strains were cultured at 37°C in Luria Broth (LB) or LB agar supplemented with the appropriate antibiotics. Kanamycin and ampicillin were used at 50 μg/mL and 100 μg/mL respectively. *P. aeruginosa* strains were cultured under agitation at 37°C in Luria Broth (LB) or LB agar.

#### Purification of pyocins and generation of antibodies

Pyocin S5 was overexpressed from *E. coli* BL21 (DE3) carrying the plasmid pPyoS5 with initial purification using a cation exchange column (CM16/10 FF, GE Healthcare; buffers: sodium phosphate 50 mM pH 6; sodium phosphate 50 mM pH 6 + NaCl 500mM). Remaining contaminants were removed using an anion exchange column (DEAE 16/10 FF, GE Healthcare; buffers: Tris 20 mM pH 8; Tris 20 mM pH 8 + 500 mM NaCl). The pyocin AP41-ImAP41 complex was purified by nickel affinity chromatography and gel filtration as described previously^23^. For both pyocins, contaminating endotoxins were removed, prior to first storage at −80°C, using endotoxin removal spin columns (Thermo Scientific #88274). LPS concentration was then verified using the Pierce LAL chromogenic endotoxin quantitation kit (Thermo Scientific 88282). Purified S5 was considered LPS-free when the levels detected were lower than that of the lowest standard of the kit at 0.1EU/mL.

#### Pyocin sensitivity assay

100 μL of *P. aeruginosa* strain culture at OD600 = 0.6 was added to 10 mL of 0.8% sterile agar and poured on an LB agar plate. Unless otherwise stated, 5 μL of pyocin, organ homogenate or serum was spotted onto overlay plates and the plates incubated overnight at 37°C.

#### In vitro killing assay in liquid cultures

*P. aeruginosa* strains were grown until OD600 = 0.6, centrifuged and resuspended in the appropriate media (LB; LB + blood; LB + 2.2’ bipyridyl) at 108 CFU/mL. An inoculum was diluted and plated for CFU counts and pyocins at various concentrations were added at t=0. Samples were then incubated at 37°C and at each time point, 40 μL were taken out and incubated for 10 minutes at room temperature with trypsin (final concentration 100 μg/mL) before being serial-diluted and plated for CFU counts.

#### Isolation of a high-level ciprofloxacin-resistant mutant

*P. aeruginosa* strain P7 was grown until OD600 = 0.6. Ten times serial dilutions of this culture were then plated on LB agar plates supplemented with ciprofloxacin 1μg/mL (Sigma) and the plates incubated overnight at 37°C. Isolated colonies were then subject to phenotypic characterization by overlay spot sensitivity assay and the minimum inhibitory concentration (MIC) was determined by standard methods as below.

#### Antibiotic sensitivity assays: minimum inhibitory concentration

The minimum inhibitory concentration (MIC) of antibiotic required to inhibit the visible growth of *P. aeruginosa* isolates was determined using a standard method^24^. The MIC assay was carried out using serial dilutions in a 96 well plate with two-fold dilutions.

#### Immunoblots

Proteins were separated by SDS-PAGE and then transferred on PVDF membranes using the Trans-Blot Turbo blotting system (Biorad). Pyocins S5 and AP41 were detected using rabbit specific polyclonal affinity-purified ɑ-S5 and ɑ-AP41 antibodies, horseradish peroxidase-coupled anti-rabbit secondary antibodies (Thermo Scientific #31460) and Clarity Western ECL reagent (Biorad). Pictures were taken using a Chemidoc system (Biorad).

#### Pyocin quantification by ELISA

Microtiter 96-well plates (Corning #9018) were coated overnight with murine sera samples diluted 1/10 in PBS. After saturation with 2% bovine serum albumin (BSA, Sigma) in PBS and 4 washes using PBS + tween 20 0.05%, adherent S5 or AP41 were detected using specific primary antibodies (rabbit polyclonal affinity-purified ɑ-S5 or ɑ-AP41 antibodies, dilution 1/1000, 2 hours incubation) and biotin-SP-conjugated AffiniPure donkey anti-rabbit IgG (Jackson ImmunoResearch Laboratories 711-065-152, dilution 1/20000, 1 hour incubation), followed by incubation with peroxidase-conjugated streptavidin (Jackson ImmunoResearch Laboratories 016-030-084, 1μg/mL in PBS, 45 min incubation) with 4 washes between each step. After extensive washes, the plates were revealed using a citrate solution containing o-phenylenediamine dihydrochloride (OPD, Thermo Scientific 34006, 10 min incubation in the dark). The reaction was stopped using 2N H2SO4 and the plate read at OD 450 nm.

#### Cytokine quantification

Serum cytokines were quantified using the Bio-plex Pro Mouse Cytokine group 1 panel 8-plex kit (Biorad M60000007A). Samples were treated according to the manufacturer’s instructions and acquired on the Bio-Plex® 200 system (Biorad).

#### Galleria mellonella larvae infection model

*G. mellonella* larvae were obtained from Livefood UK, kept in darkness at room temperature and were used up to one week following arrival. Healthy larvae with no melanisation were used for all experiments. *P. aeruginosa* was grown under agitation in LB broth at 37°C to an OD_600_ = 0.6. Cells were then washed twice in sterile PBS and diluted to the desired inoculum in PBS. Inoculums were serially diluted and plated on LB agar plates just before administration for CFU counting. Groups of 10 larvae were injected with 10 μL of bacterial suspension in the hemocoel via the last right pro-limb. Following challenge, larvae were placed in an incubator at 37°C. Larvae were treated 3 hours post-infection by injection of 10 μL of BSA, pyocin or ciprofloxacin in PBS at various concentrations in the hemocoel via the last left pro-limb. Survival was followed for 48 hours, larvae were considered dead when unresponsive to touch. For each experiment, groups of uninfected larvae (n=10) were injected with PBS, BSA, pyocin or antibiotics as a negative control. No deaths were recorded for these control groups.

#### Stability and clearance of pyocins in vivo

LPS-free and sterile filtered pyocin S5 or pyocin AP41 in PBS were injected intravenously (IV) at a concentration of 1mg/mL in 8-10 weeks old female Balb/c mice (Charles River). Mice were culled at different time points following the injection (earliest 10 minutes, latest 33 hours), blood was obtained via cardiac puncture immediately following carbon dioxide asphyxiation and organs were sampled. The blood was left to coagulate for 1 hour at room temperature before being centrifuged for 10 min at 3000 rpm to obtain the serum. Treatment of organs was as followed: PBS supplemented with protease inhibitor (1 tablet cOmplete protease inhibitor, Roche in 35 mL of PBS) was added to the frozen organs (volume depending on the weight of the organ) before homogenization using a handheld homogenizer (Omni). Homogenized organs were then centrifuged at 4°C for 20 min at 13000 rpm and supernatant stored with the sera at −80°C until analysis by pyocin sensitivity assay and immunoblots.

#### Murine sepsis model

7 weeks old female Balb/c mice were infected by IV injection with a lethal dose of exponentially growing *P. aeruginosa* (2 × 10^7^ CFU). Five hours post-infection, mice were treated with LPS-free, sterile filtered BSA or pyocins by IV injection. Clinical health was then monitored for up to one week and mice culled when reaching the endpoint either when the clinical score set as threshold (severity score =7, table S1) was reached or at the pre-defined endpoint of the experiment. Blood (with or without heparin) and organs were then collected. For CFU counts, blood was serial-diluted and plated immediately on LB agar. Organs were homogenized in PBS using a hand-held homogenizer (Omni) and plated after serial dilutions. Sera was stored at −80°C until analysis.

#### Ethics statement

All animal experiments were performed in accordance with the UK Animals (Scientific procedures) Act, authorized under a UK Home Office License and all procedures were approved by the animal project review committee of the University of Glasgow. The project license number assigned by the animal project review committee of the University of Glasgow was P079A1B53.

## Results

### Pyocins S5 and AP41 are bactericidal in different conditions in vitro

First, we determined the ability of pyocins S5 and AP41 to kill *P. aeruginosa* strains under different conditions in vitro. Using the pyocin sensitivity spot assay showed that the P7 clinical isolate of *P. aeruginosa* was sensitive to both S5 and AP41, with S5 being more potent (Minimum effective dose (MED) = 0.1 ng than AP41 (MED = 2 ng) (Figure 1A). In LB liquid culture, both pyocins had bactericidal activity, causing a sharp decrease in CFU counts after 30 min incubation with S5 and after one hour incubation with AP41 (Figure 1B). Interestingly, while S5 had a more rapid effect than AP41, CFU counts started increasing again after 2 hours of incubation with S5, which was not the case with AP41. When a combination of both pyocins was used, a combination of both phenotypes was observed, with a quick but stable decrease of CFU counts over time (Figure 1B). As the pore-forming pyocin S5 has been shown to use the ferripyochelin receptor FptA^17^, activity of both pyocins was tested in LB supplemented with the iron chelator 2.2’-bipyridyl at a final concentration of 2 mM. As expected, the addition of 2.2’ bipyridyl significantly increased the bactericidal activity of S5 (Figure 1C). However, the addition of the iron chelator had no effect on AP41 activity suggesting that its receptor, the identity of which is not known, is unlikely to be involved in iron uptake. In a similar way, to determine if pyocins retain their bactericidal activity in blood, bacterial counts were followed during incubation with recombinant pyocins in LB supplemented with 50% fresh murine blood (Figure 1C). The presence of blood increased pyocin S5 activity with a 10-fold decrease in CFU numbers compared to incubation in LB alone. Incubation with AP41 in 50% fresh blood decreased its antimicrobial activity; however, this decrease was not significant (Figure 1C). These results show that both pyocins retain their activity in blood.

**Figure 1:**
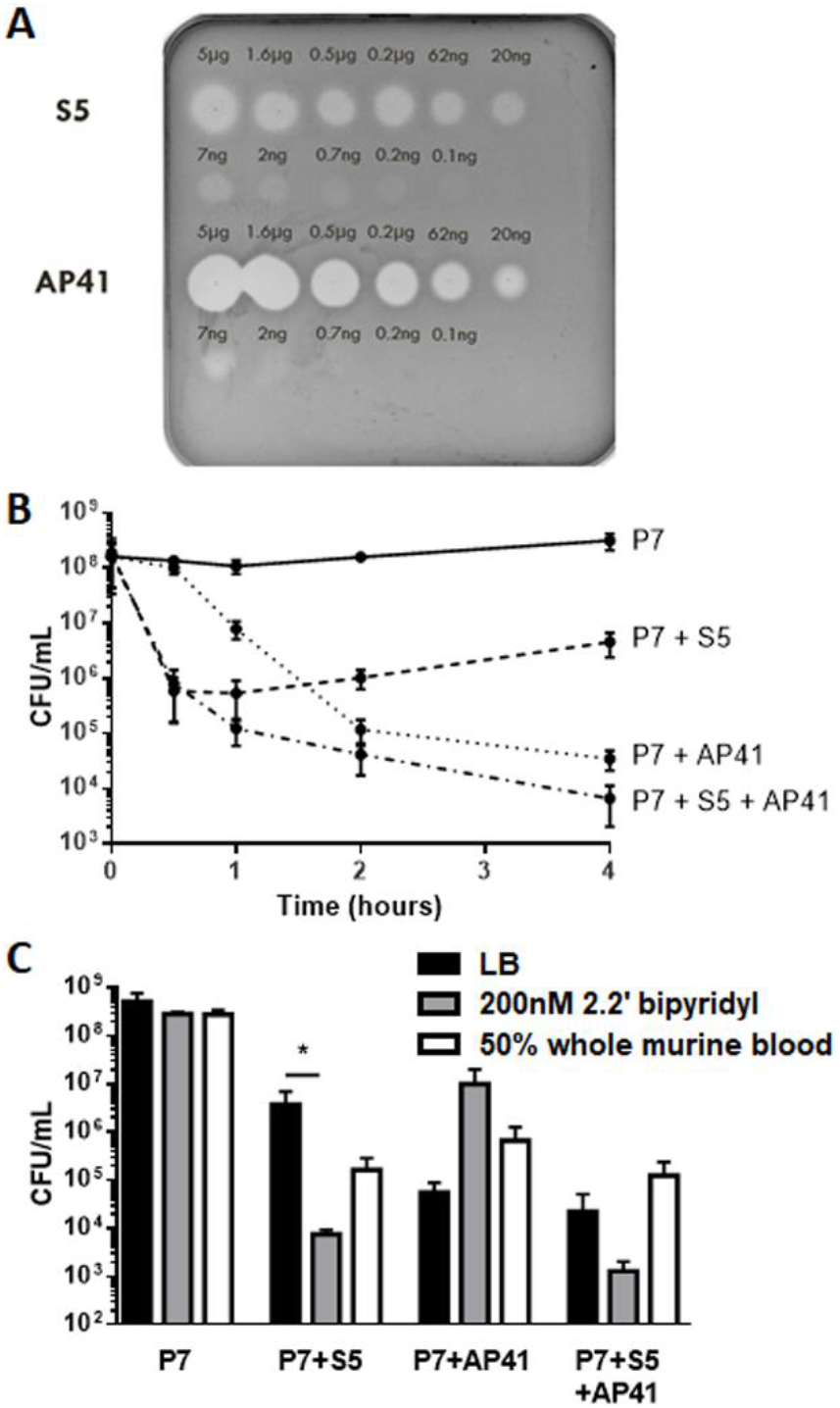
Purified pyocins S5 and AP41 are active in vitro. **(A)** Spot tests to determine the activity of pyocins S5 and AP41 against *P. aeruginosa* strain P7. 5 μL of a range of concentration of pyocin S5 / AP41 were spotted on a growing lawn of *P. aeruginosa* Presence of clear zones indicate pyocin activity. **(B)** Bacterial counts over time of *P. aeruginosa* incubated in LB media supplemented with PBS (control) or 10 μg/mL of S5, AP41 or S5 and AP41. **(C)** Bacterial counts of *P. aeruginosa* after 4 h of incubation with PBS, S5, AP41 or S5 + AP41 (10 μg/mL) in different conditions: LB, LB + 200 nM 2.2’ bipyridyl or LB + fresh whole murine blood (50%). Columns represent the average CFU counts in CFU/mL from three experiments. Error bars represent the SD and asterisks indicate significant differences as assessed by one-way ANOVA (* P<0.05).

#### Pyocins S5 and AP41 are able to treat lethal *P. aeruginosa* infection in *Galleria mellonella* larvae

To determine if pyocins AP41 and S5 were able to treat a lethal systemic infection in vivo, *Galleria mellonella* larvae were infected with 5 × 10^3^ CFU of *P. aeruginosa* P7 clinical isolate and treated 3 hours following infection with different doses of pyocins. Larvae were monitored, and survival assessed 24 hours post-infection (Figure 2A). While most untreated larvae were dead by 24 hours post-infection, we observed 80-100% survival of larvae treated with either S5 (10 ng to 10 μg), or AP41 (100 ng to 10 μg), showing the effectiveness of both pyocins to protect against a lethal *P. aeruginosa* infection in this model. At the lowest treatment dose tested, S5 was able to rescue larvae survival, but there was no increase in survival compared to the untreated larvae when using AP41 (Figure 2A).

**Figure 2:**
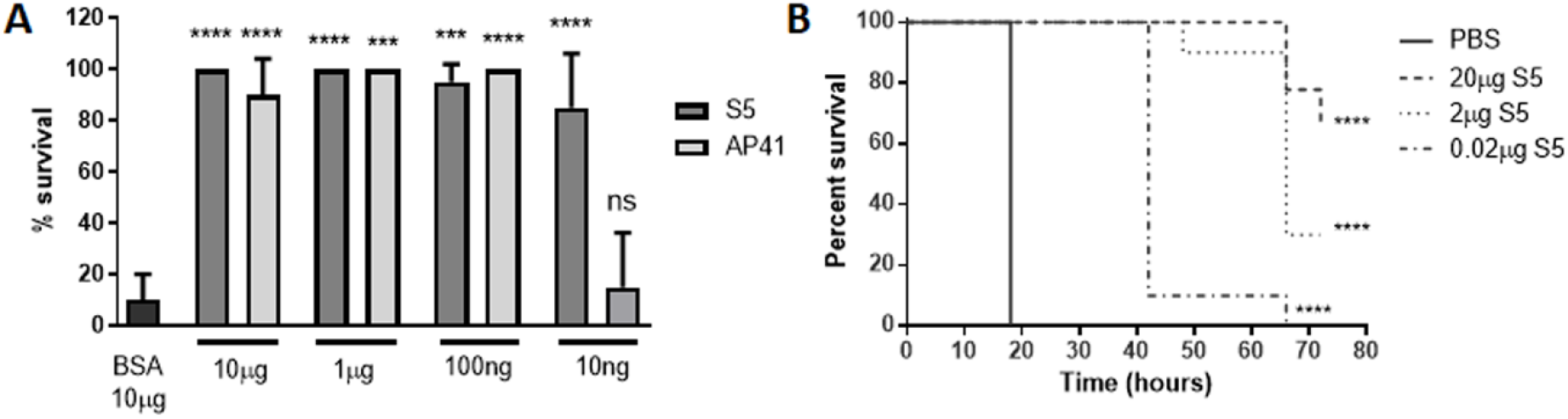
Purified pyocins S5 and AP41 are able to treat *Pseudomonas aeruginosa* infected *Galleria mellonella* larvae. **(A)** 24 h survival of larvae infected with *P. aeruginosa* P7 strain and treated 3 hours post-infection with BSA, S5 or AP41. Columns represent the mean survival from two experiments (n=10 larvae per group per experiment). Error bars represent the SD. Asterisks indicate significant differences relative to larvae treated with PBS, as assessed by one-way ANOVA (ns, non significant; ****, P<0.0001; ***, P<0.001). **(B)** Kaplan-Meier survival curves of larvae infected with *P. aeruginosa* P7 strain and treated 3 hours post-infection with different doses of S5. Survival curves are representative of two distinct experiments. Asterisks indicate significant differences relative to larvae treated with PBS, as assessed by the log-rank (Mantel-Cox) test (****P<0.0001).

In a separate experiment extending to 5 days post-infection, showed that following S5 treatment survival decreased over time even at the highest treatment dose (Figure 2B). When AP41 was used as treatment instead of S5, a 100% mortality rate was observed 48 hours post-infection even for larvae treated at the highest dose (data not shown). In both cases, S5 and AP41 were able to delay larval death, which may reflect that the pyocins are cleared or degraded before total clearance of the bacteria, or that infection has reduced the larval lifespan. However, the differences in the survival following pyocin treatment were all significantly different from larvae treated with BSA alone.

### Pyocin S5 is able to treat an antibiotic-resistant Pseudomonas infection in the *Galleria larvae* model

Eradication of *P. aeruginosa* is becoming increasingly difficult because of the increasing prevalence of drug-resistant isolates. Ciprofloxacin is one of the drugs commonly used to treat *P. aeruginosa*; however, high-level resistance can be acquired rapidly, rendering the drug ineffective. In order to test the potency of pyocin S5 to treat an infection caused by a high-level antibiotic-resistant strain of *P. aeruginosa*, we generated a high-level ciprofloxacin-resistant mutant (P7ciproR+) derived from the clinical isolate P7. The initial P7 isolate is already resistant to ciprofloxacin (MIC = 2 μg/mL); the derived high-level resistant mutant had a MIC of 16 μg/mL. In vitro, this mutant retained its susceptibility to pyocin S5 (Figure 3A). The efficacy of S5 against the P7ciproR+ mutant was assessed in vivo using the Galleria larvae model of systemic infection (Figure 3B-C). Representative curves are shown in Figure 3B and C with additional replicates in Supplementary Figure 1. As expected, while survival of larvae infected with the WT P7 strain was dramatically increased by treatment with ciprofloxacin (10 μg, 80% survival 48 hours post-infection), the same treatment was only able to delay the deaths of the larvae infected with the P7ciproR+ mutant, with 100% mortality observed 27 hours post-infection. S5 was able to rescue the larvae infected with the WT P7 strain or P7ciproR+ mutant equally effectively (90% survival 48 hours post-infection for both), demonstrating that pyocin S5 is able to treat a highly antibiotic-resistant *P. aeruginosa* infection in this Galleria larvae model.

**Figure 3:**
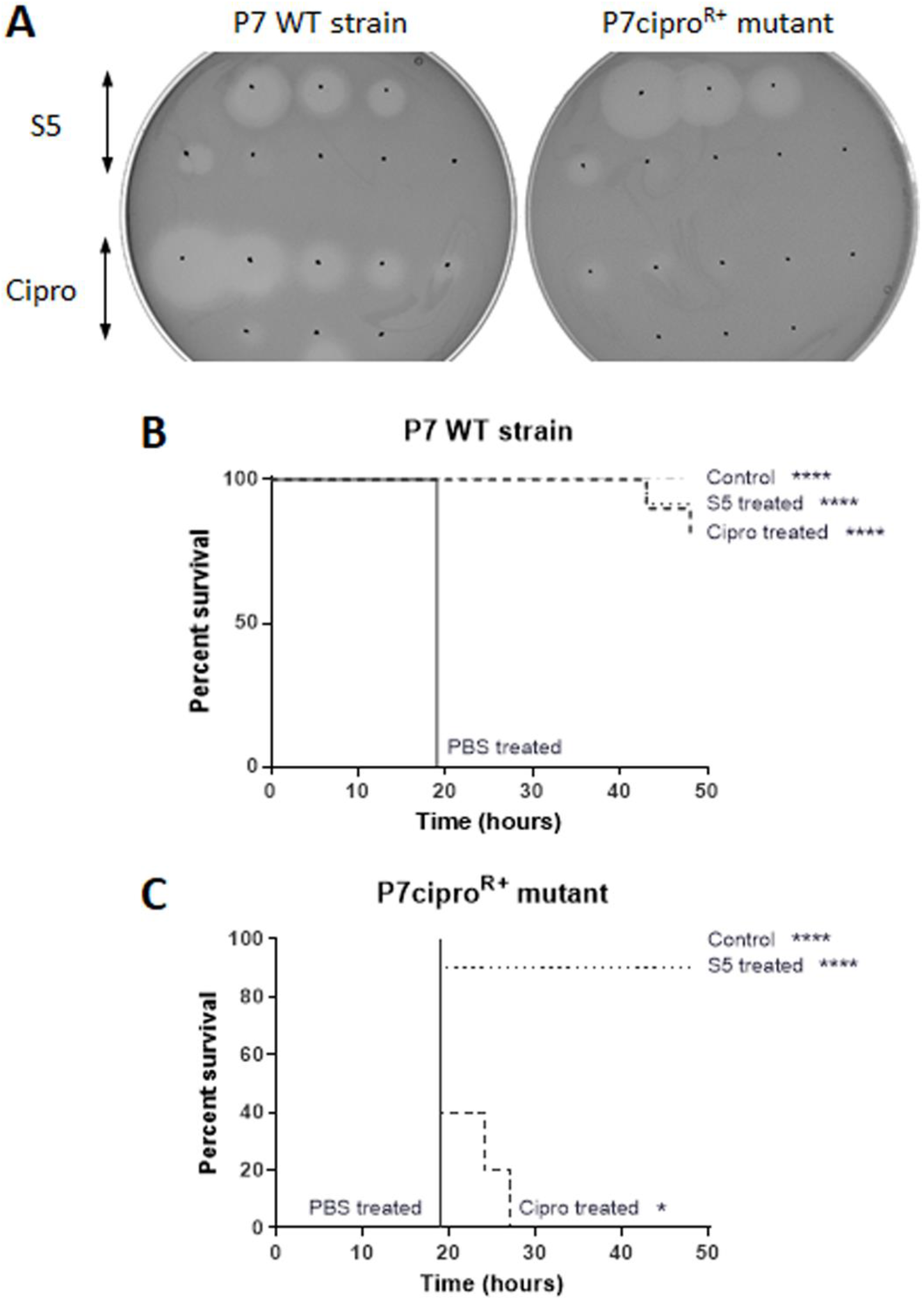
Purified pyocin S5 is active against ciprofloxacin-resistant *Pseudomonas aeruginosa* in vitro and in vivo. **(A)** Spot tests to determine the activity of ciprofloxacin and pyocin S5 against *P. aeruginosa* strain P7 and P7-derived ciprofloxacin-resistant mutant. 3 μL of a range of concentration of ciprofloxacin (3 μg – 23 ng, 2X dilutions) and pyocin S5 (3 μg – 0.3 pg, 10X dilutions) were spotted on a growing lawn of *P. aeruginosa*. The presence of clear zones indicates pyocin or antibiotic activity. **(B-C)** Kaplan-Meier survival curves of larvae infected with the WT *P. aeruginosa* P7 strain **(B)** or the P7-derived ciprofloxacin-resistant mutant (P7ciproR+) **(C)** (n=10 per group) treated 3 hours post-infection with ciprofloxacin (10 μg) or pyocin S5 (5 μg). Control uninfected larvae (n=10 per group) were injected with PBS. Groups of uninfected larvae (n=10 per group) were also injected with ciprofloxacin (10 μg) or pyocin S5 (5 μg) and all survived but are not represented here. Survival curves are representative of two distinct experiments. Asterisks indicate significant differences relative to larvae treated with PBS, as assessed by the log-rank (Mantel-Cox) test (* P<0.05; ****P<0.0001).

### Dissemination, degradation and activity in vivo in mice

The in vitro and in vivo studies show that pyocins AP41 and S5 are active in the presence of fresh blood and retain activity in an invertebrate infection model. To begin exploring the utility of pyocins in treating *P. aeruginosa* blood infections, we first investigated their distribution, clearance and stability in vivo. High doses of AP41 or S5 (200 μg) were injected intravenously into mice and the presence and activity of the pyocins in different organs was measured at different time points following infection. The concentrations of AP41 and S5 in serum over time were determined by ELISA using polyclonal specific AP41 or S5 antibodies, respectively (Figure 4A). The serum concentration of S5 decreased sharply almost immediately following injection, being detectable in the serum only up to 1 hour post-injection. By contrast, the serum concentration of AP41 only slightly decreased over time. Dissemination of S5 or AP41 from the blood to different organs was then determined by

**Figure 4:**
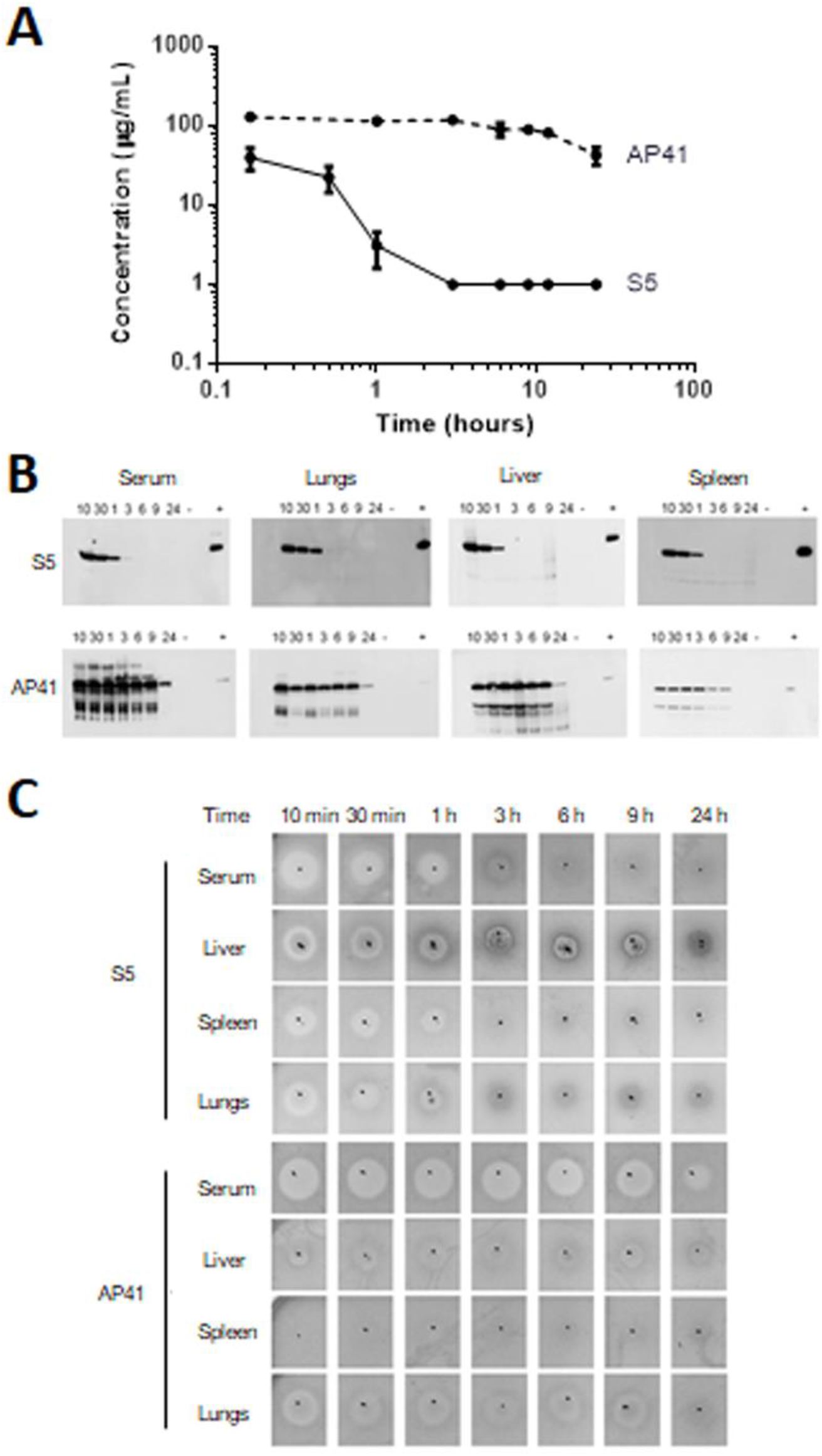
Pyocins S5 and AP41 have different dissemination and clearance profiles in mice. **(A)** Serum concentration of S5 and AP41 determined by ELISA following IV injection of 200 μg of pyocins in mice. Results represent the mean from three independent experiments, two mice per time point and per experiment. Error bars represent the SD. **(B)** Detection of pyocins S5 and AP41 in serum and organ homogenates at different time-points (10 min, 30 min, 1 h, 3 h, 6 h, 9 h, 24 h) following injection using S5- or AP41-specific affinity purified antibodies. Positive control (+) = 1.25 ng of purified S5 or AP41. Negative control (−) = Corresponding homogenized organ from a mouse that did not receive injections of S5 or AP41. **(C)** Detection of pyocin S5 or AP41 activity in serum or homogenized organs at different time-points following IV injection. 5 μL of homogenized organ or serum were spotted on a growing lawn of *P. aeruginosa* P7 strain. Presence of clear zones indicate pyocin activity.

Western blot (Figure 4B). Pyocin S5 was detected up to 1 hour following injection in serum, confirming the ELISA results, as well as in spleen, lungs and liver showing good distribution of the pyocin within the mouse. Surprisingly, a unique full-sized S5 band was observed with the absence of degradation products, even at later time-points. Pyocin AP41 was detected up to 24 hours post-injection in serum, spleen, lungs and liver, showing good distribution and stability. Finally, we tested the activity of both pyocins from the same samples by sensitivity assay (Figure 4C). As expected, zones of inhibition were detected up to one hour post-injection in serum, lungs and spleen of S5-treated mice, and up to 24 hours in serum and lungs of AP41-treated mice. The anti-bacterial activity shown through the sensitivity assay correlates well with results from both western blot and ELISA techniques, other than a lack of bactericidal activity in the spleens of AP41 treated mice. Overall, these results show that both pyocin S5 and AP41 are able to disseminate quickly through the organism, with S5 being cleared faster than AP41.

### IV injection of S5 and AP41 does not induce an inflammatory response

To determine if injection of pyocins on their own induce a pro- or anti-inflammatory response in non-infected mice when injected intravenously, we quantified levels of pro-inflammatory (IL-1β and TNF-ɑ) and anti-inflammatory (IL-10) cytokines in serum samples from S5- and AP41-treated mice and compared these to the levels observed in non-injected mice. Neither pyocin S5 nor pyocin AP41 induced production of these cytokines, with levels in treated animals remaining very similar to those of control mice (Supplementary Figure 2).

### Pyocins S5 and AP41 improve survival of septic mice

To determine if pyocins S5 and AP41 are able to improve survival rates of septic mice when administered post-infection, 7 week-old mice were infected by intravenous injection of a lethal dose of P7 strain and treated by intravenous injection of 2 μg of pyocin S5 or AP41 5 hours post-infection. Representative experiments are shown in Figure 5 Both pyocins S5 and AP41 were able to increase survival in this model from 33% to 83% and 67%, respectively (Figure 5A); this is significant for S5 but not AP41. A second injection of 2 μg of S5 did not improve the survival rate, nor did the use of a cocktail of S5 and AP41. To determine if increasing the concentration would result in increased survival of infected mice, different doses of pyocin S5 were used (200 μg, 20 μg, 2 μg) and the mice monitored for 72 hours. Treatment with 2 μg, 20 μg or 200 μg of pyocin S5 all had a very similar effect on survival (Figure 5B).

**Figure 5:**
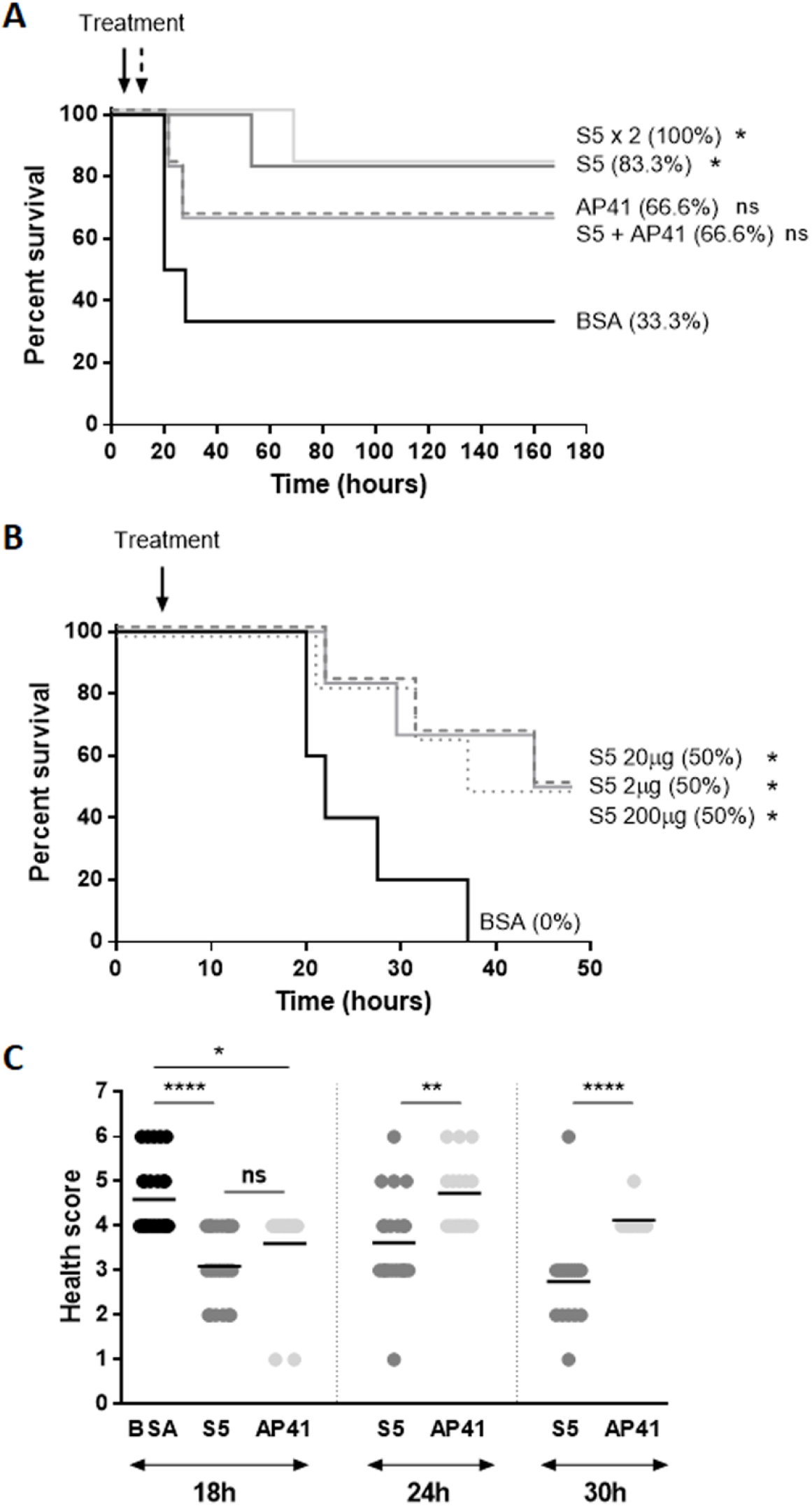
Pyocins S5 and AP41 improve survival of septic mice. **(A)** Kaplan-Meier survival curves of mice (n=6 per group) infected by IV with 5.6 × 10^7^ CFU of *P. aeruginosa* strain P7 and treated by IV injection 5 h post-infection with a single injection of 2 μg of BSA, S5 or AP41, with an injection of a cocktail of S5+AP41 (2 μg) or treated by two injections of 2 μg of S5 (5 and 11 hours post-infection). **(B)** Kaplan-Meier survival curves of mice infected by IV with 4 × 10^7^ CFU of *P. aeruginosa* strain P7 and treated by IV injection 5h post-infection with 200 μg (dotted line), 20 μg (dashed line), 2 μg (solid line) of S5 or 200 μg of BSA. Survival curves are representative of at least two independent experiments. Asterisks indicate significant differences relative to mice treated with BSA, as assessed by the log-rank (Mantel-Cox) test (ns: non-significant; *: P<0.05). **(C)** Clinical scores of infected mice 18h, 24h and 30h post-infection. Data was collected from four different experiments. Asterisks indicate significant differences as assessed by one-way ANOVA (ns: non-significant; *: P<0.05; **: P<0.01; ****: P<0.0001).

Although there was an increase of survival for the mice treated with pyocin S5 or AP41, they still displayed severe signs of infection. Following infection, mice were monitored regularly and a health score given to each of them using a scoring system for disease severity (Table S1, Figure 5C). The onset of the disease for sham-treated mice occurred between 18 and 24 hours post-infection. At 18 hours post-infection mice were given an average score of 4.6, 3.0 and 3.6 for sham-treated, S5-treated and AP41-treated mice respectively, a significant reduction at that time point for the scores of pyocin-treated mice. As most sham-treated mice did not survive beyond the 18 hours post-infection time point, no clinical scores were included for that group after this time-point. At 24 hours post-infection, average health scores increased to 3.6 and 4.7 for S5-treated and AP41-treated mice respectively, suggesting that pyocins produce a delay in disease progression. Comparison of the evolution of health scores between S5- and AP41-treated mice 24 hours and 30 hours post-infection show significantly lower health scores and a more rapid recovery of S5-treated mice. The mice were monitored for up to 7 days following infection and once completely recovered, there was no relapse observed and no delayed murine deaths.

To determine if pyocins S5 and AP41 are able to reduce bacterial load in mice following infection, groups of 6 mice were culled 6 hours, 12 hours and 19 hours post-infection and bacterial counts in blood, liver, lungs and spleen from treated mice were compared to that of sham-treated mice (Figure 6). 19 hours was set as the last time point as by that time the first of infected mice reach the severity limit set-up in the protocol. In these experiments, S5 significantly reduced bacterial load in the blood at 6 hours and 12 hours post-infection. However, while there was a trend towards a reduction of bacterial numbers in the blood of AP41-treated mice, this was not significant (Figure 6A), correlating with the greater survival of S5-treated mice compared to AP41-treated mice. As both bacteria and pyocins will disseminate from the blood to organs, we also determined bacterial load in organs of untreated and treated mice. There was no significant reduction of bacterial numbers in liver and lungs of pyocin-treated mice (Figure 6B-C). In the spleen, we observed a significant decrease of CFU counts 19 hours post-infection for AP41-treated mice (Figure 6D). Bacterial counts in the remaining healthy mice at the end of experiments were also investigated. No bacteria were found in the blood of mice that recovered from infection after being treated with 2 μg of S5 (n=6, t=72h post-infection; n=5, t=168 h post-infection) or with 2 μg of AP41 (n=6, t=72 h post-infection; n=5, t=168 h post-infection).

**Figure 6:**
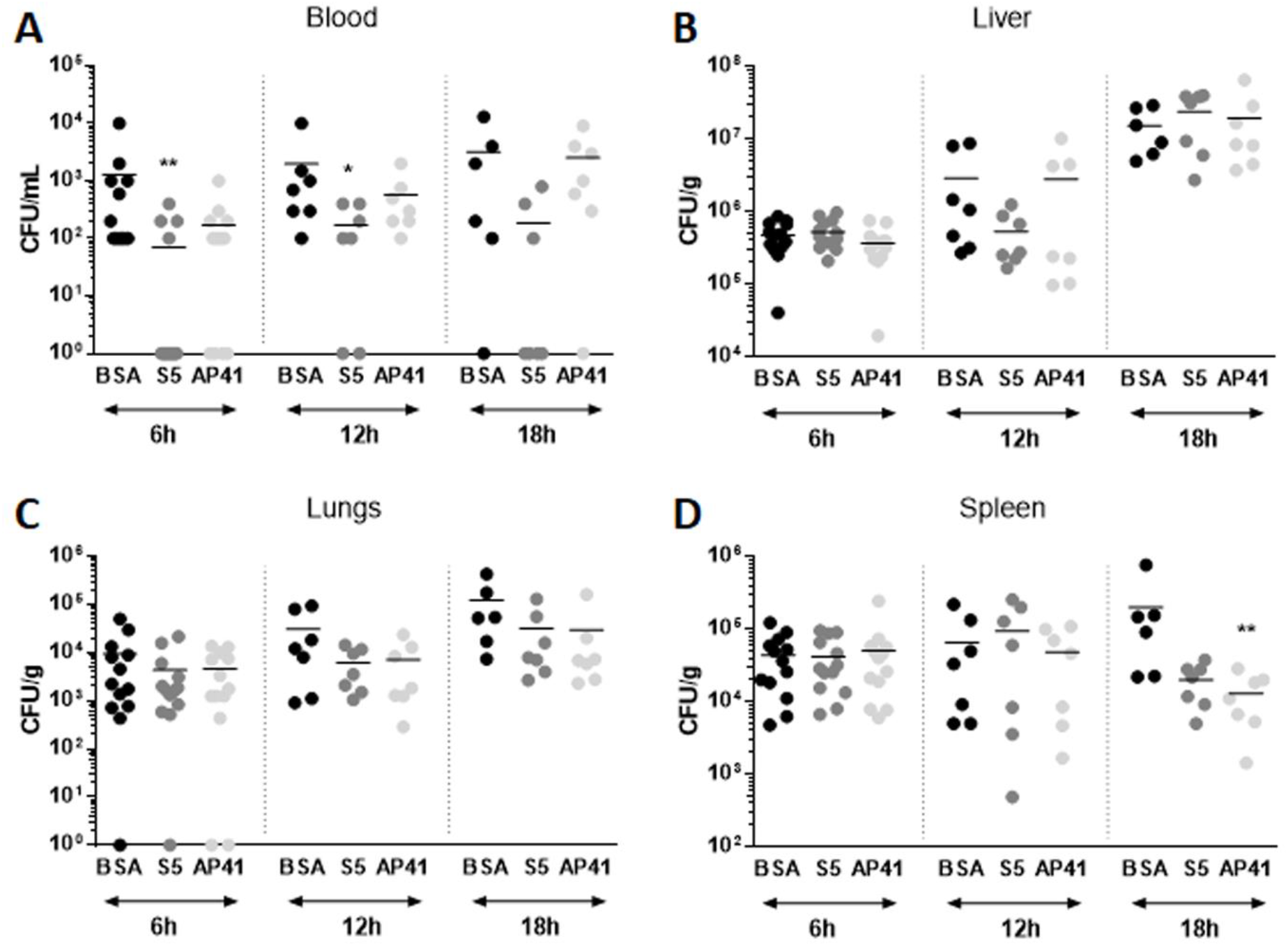
Pyocins S5 and AP41 improve survival of septic mice. Bacterial counts 6h, 12h and 18 h post-infection in the blood **(A)**, liver **(B)**, lungs **(C)**, spleen **(D)** of infected mice treated with 2 μg of BSA, 2 μg of S5 or 2 μg of AP41. Asterisks indicate significant differences relative to sham-treated mice, as assessed by the Kruskal-Wallis test followed by Dunn’s multiple comparisons test (* P<0.05; ** P<0.01; ***P<0.0001). When not indicated, the differences were not statistically different.

### Pyocins S5 and AP41 do not affect cytokine production

The severity of sepsis is linked to the host response to the infection, as sepsis is characterized by a massive and auto-amplifying cytokine production know as a “cytokine storm”. To determine if pyocin treatment was able to affect cytokine production, the concentrations of serum pro- and anti-inflammatory cytokines (IL-1β, IFN-γ, TNF-ɑ and IL-10) were quantified for each mouse before infection, after infection and after pyocin injection. As expected, cytokine levels increase significantly as a response to bacterial infection. However, there was no significant difference in cytokine levels following treatment of infected mice by BSA, pyocin S5 or pyocin AP41, suggesting that S5 and AP41 treatments do not have a direct immunomodulatory effect in this model (Figure 7A-D).

**Figure 7:**
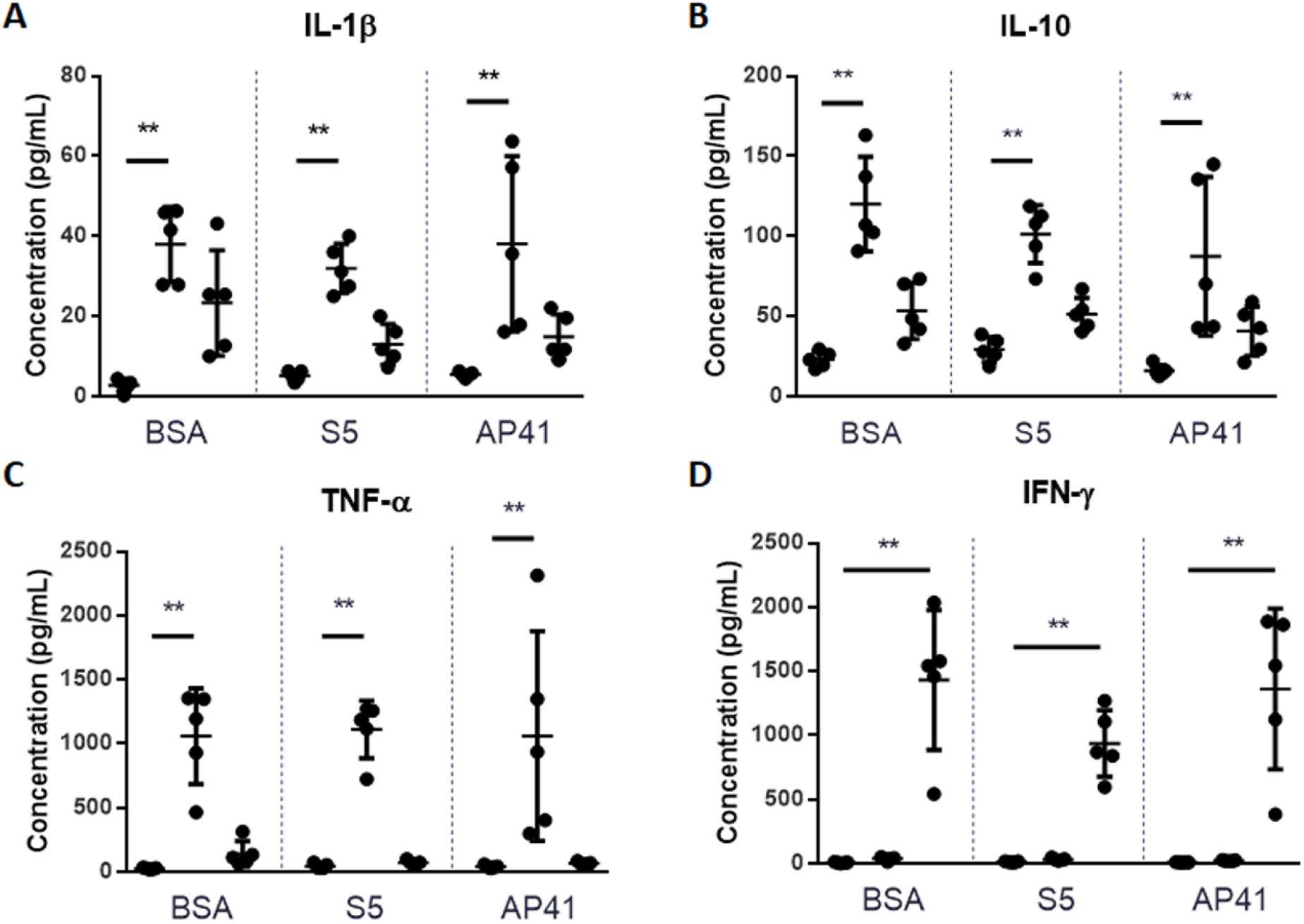
Treatment of septic mice with pyocin S5 or AP41 does not alter cytokine production. Serum concentration of IL-1β **(A)**, IL-10 **(B)**, TNF-ɑ **(C)** and IFN-γ **(D)** in mice one-hour post-treatment. Each symbol represents the mean value obtained per mouse. The bar shows the mean value, and the error bars the SD Asterisks indicate significant differences, as assessed by the Kruskal-Wallis test followed by Dunn’s multiple comparisons test (** P<0.01). When not indicated, the differences were not statistically different.

### In vivo pyocin treatment did not result in pyocin resistant/tolerant colonies

Previously, we have isolated one AP41-tolerant strain from AP41-treated mice in an acute respiratory infection model^23^. To determine if pyocin tolerance was acquired following intravenous treatment of septic mice, CFU recovered from mice at different time points following infection were tested for pyocin sensitivity (n = 119, Table S2). The colonies tested were recovered from mice treated with S5 (28.6%), AP41 (50.4%), S5 + AP41 (6.7%) or BSA as a control group (14.3%). No difference in sensitivity to pyocin S5 or AP41 was observed in colonies recovered from mice compared to that of the original P7 strain.

## Discussion

*P. aeruginosa* is one of the leading cause of sepsis with significant patient mortality and high health-care costs. The development of bacteriocins as novel species-specific protein antibiotics is a promising strategy in the face of increasing antibiotic-resistance. Previously, our group demonstrated the efficacy of pyocins to treat pneumonia in an acute murine lung infection model^23^. In this work, we aimed to test the efficacy of two pyocins for the treatment of *P. aeruginosa* sepsis. S5 and AP41 were chosen for their different uptake mechanisms, cytotoxic activities and for their activity against a broad range of clinical isolates. Not much is known about the mechanisms of entry into the target cells by pyocins. While pyocin S5 utilises the common polysaccharide antigen and the TonB-dependent iron-siderophore transporter FptA to target *P. aeruginosa*, the mechanism of pyocin AP41 uptake is currently unknown^16,17^. However, we show that there is no difference in efficacy of AP41 in vitro in presence or absence of an iron-chelating agent (2.2’ bipyridyl) suggesting that the receptor is unlikely to be an iron-siderophore receptor. Importantly, both pyocins have an in vitro bactericidal activity that they retain in presence of fresh murine blood.

In this study, single injections of two *P. aeruginosa*-specific pyocins are shown to improve the survival of septic mice. Following intravenous infection of the mice, *P. aeruginosa* is able to disseminate to different organs, highlighting the importance of the ability of the chosen anti-microbial to spread from the blood to the organs in order to target the pathogen. When injected directly in the blood, pyocins do not provoke an immune reaction, remain stable and retain their activity for a minimum of 1 hour for S5 and 24 hours for AP41. We demonstrate that both pyocins are able to disseminate into the liver, the spleen and the heart where they are active. Surprisingly, while AP41 is found to remain stable and active in the blood and organs for a longer period in vivo, and found to be more efficient for colony-counts reduction in vitro, it is actually less efficient as treatment in a murine model of sepsis. Indeed, the observed survival rate of mice treated with AP41 was lower than that of mice treated with S5. Additionally, we saw no significant reduction in CFU counts in the blood of infected mice following treatment with AP41. Interestingly, the difference in efficacy of treatment with S5 and AP41 in infected mice was also observed in infected larvae, further validating the use of *G. mellonella* larvae as a bridge between in vitro assay and murine infection models. These observations could be due to mechanistic difference between AP41 and S5 in in vivo conditions, or an interaction of AP41 with host proteins, preventing or decreasing its activity in vivo against *P. aeruginosa*. Further studies would be necessary to test these different hypotheses.

A single injection of pyocin is not sufficient to produce a 100% survival rate; moreover, the treated mice still display severe symptoms of infection, with high bacterial counts found in organs. The bacteria recovered from animals following infection and treatment were tested for sensitivity to pyocins, and no pyocin-resistant or tolerant mutants were found. This suggests that instead of completely clearing the bacterial infection, pyocins reduce numbers of viable bacteria to a level where innate immune protection can allow animals to survive. It is possible that giving a continuous intravenous infusion of pyocin will increase the survival rate. However, we show that giving a second injection of pyocin S5 12 hours post-infection did not improve survival compared a single injection. In the most recent Sepsis-3 consensus, sepsis is defined as a life-threatening organ dysfunction caused by a dysregulated host-response to infection^25^. An ideal treatment against sepsis would be the use of a cocktail of antimicrobials and immunomodulatory agents, targeting the pathogen while also restoring immune homeostasis. While pyocins S5 and AP41 have a bactericidal activity, they have no effect on the production of the pro- and anti-inflammatory cytokines tested (IL-1β, IL-10, IFN-γ, TNF-ɑ).

Finally, with the rise of antimicrobial resistance being an increasingly worrying problem, this study shows the potential of pyocins as targeted protein antimicrobials against antibiotic-resistant strains, as pyocin S5 is able to protect *G. mellonella* larvae against a lethal high-level ciprofloxacin-resistant *P. aeruginosa* challenge.

## Transparency Declaration

The University of Glasgow has filed a patent on the use of pyocins to treat *Pseudomonas aeruginosa* lung infection with D.W as an inventor. There are no other conflicts of interest for any of the authors.

## Supplementary Figures

**Supplementary Figure 1:**
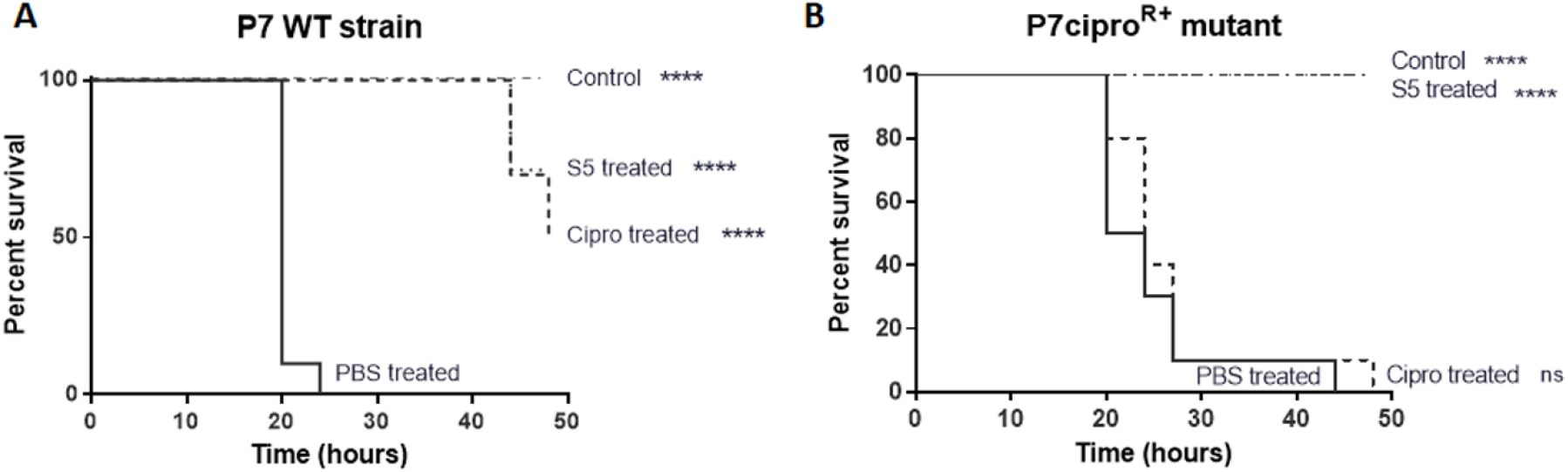
Purified pyocin S5 is efficient against high-level ciprofloxacin-resistant *Pseudomonas aeruginosa* in vivo. Kaplan-Meier survival curves of larvae infected with the WT *P. aeruginosa* P7 strain **(A)** or the P7-derived ciprofloxacin-resistant mutant (P7ciproR+) **(B)** (n=10 per group) treated 3 h post-infection with ciprofloxacin (10 μg) or pyocin S5 (5 μg). Control uninfected larvae (n=10 per group) were injected with PBS. Groups of uninfected larvae (n=10 per group) were also injected with ciprofloxacin (10 μg) or pyocin S5 (5 μg) and all survived but are not represented here. Survival curves are representative of two distinct experiments. Asterisks indicate significant differences relative to larvae treated with PBS, as assessed by the log-rank (Mantel-Cox) test (ns: non-significant; ****P<0.0001).

**Supplementary Figure 2:**
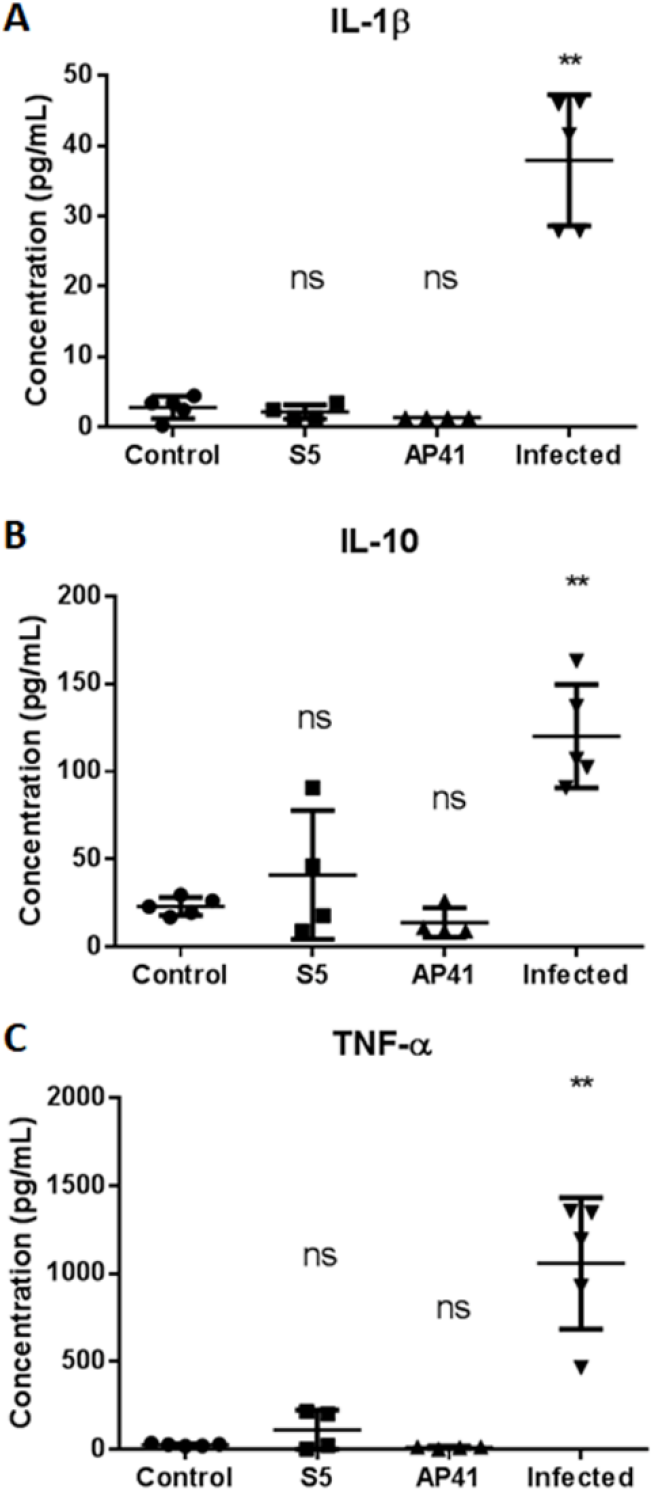
Effect of Pyocins S5 and AP41 on cytokine production. Serum levels of cytokines were quantified from mice having received an IV injection of S5 or AP41 (200 μg, 1 h post-injection), from un-injected healthy mice (negative control) and from mice infected by IV with *P. aeruginosa* P7 strain (2 h post-infection). Data points represent levels of IL-1β (A), IL-10 (B) and TNF-ɑ (C) obtained for each mouse; the bar shows the mean value, and the error bars the SD. Asterisks indicate significant differences as assessed by one-way ANOVA (ns: non-significant; **: P<0.01).

## References

1. Kang, C.-I. et al. Bloodstream infections caused by antibiotic-resistant gram-negative bacilli: risk factors for mortality and impact of inappropriate initial antimicrobial therapy on outcome. Antimicrob. Agents Chemother. 49, 760–6 (2005).

2. Vidal, F. et al. Epidemiology and outcome of Pseudomonas aeruginosa bacteremia, with special emphasis on the influence of antibiotic treatment. Analysis of 189 episodes. Arch. Intern. Med. 156, 2121–6 (1996).

3. Nathwani, D., Raman, G., Sulham, K., Gavaghan, M. & Menon, V. Clinical and economic consequences of hospital-acquired resistant and multidrug-resistant Pseudomonas aeruginosa infections: A systematic review and meta-analysis. Antimicrob. Resist. Infect. Control 3, 32 (2014).

4. Johnson, L. E. et al. Pseudomonas aeruginosa bacteremia over a 10-year period: multidrug resistance and outcomes in transplant recipients. Transpl. Infect. Dis. 11, 227–34 (2009).

5. Suárez, C. et al. Influence of carbapenem resistance on mortality and the dynamics of mortality in Pseudomonas aeruginosa bloodstream infection. Int. J. Infect. Dis. 14 Suppl 3, e73–8 (2010).

6. Morata, L. et al. Influence of multidrug resistance and appropriate empirical therapy on the 30-day mortality rate of Pseudomonas aeruginosa bacteremia. Antimicrob. Agents Chemother. 56, 4833–7 (2012).

7. Hattemer, A. et al. Bacterial and clinical characteristics of health care- and community-acquired bloodstream infections due to Pseudomonas aeruginosa. Antimicrob. Agents Chemother. 57, 3969–75 (2013).

8. Livermore, D. M. Multiple Mechanisms of Antimicrobial Resistance in Pseudomonas aeruginosa: Our Worst Nightmare? Clin. Infect. Dis. 34, 634–640 (2002).

9. Carmeli, Y., Troillet, N., Eliopoulos, G. M. & Samore, M. H. Emergence of antibiotic-resistant Pseudomonas aeruginosa: Comparison of risks associated with different antipseudomonal agents. Antimicrob. Agents Chemother. 43, 1379–1382 (1999).

10. WHO | Prioritization of pathogens to guide discovery, research and development of new antibiotics for drug resistant bacterial infections, including tuberculosis. WHO (2017).

11. Surveillance of antimicrobial resistance in Europe 2018. https://www.ecdc.europa.eu/en/publications-data/surveillance-antimicrobial-resistance-europe-2018.

12. Behrens, H. M., Six, A., Walker, D. & Kleanthous, C. The therapeutic potential of bacteriocins as protein antibiotics. Emerg. Top. Life Sci. 1, 65–74 (2017).

13. Hammami, R., Fernandez, B., Lacroix, C. & Fliss, I. Anti-infective properties of bacteriocins: an update. Cell. Mol. Life Sci. 70, 2947–67 (2013).

14. Sharp, C., Bray, J., Housden, N. G., Maiden, M. C. J. & Kleanthous, C. Diversity and distribution of nuclease bacteriocins in bacterial genomes revealed using Hidden Markov Models. PLoS Comput. Biol. 13, e1005652 (2017).

15. Ghequire, M. G. K. & De Mot, R. Ribosomally encoded antibacterial proteins and peptides from Pseudomonas. FEMS Microbiology Reviews vol. 38 523–568 (2014).

16. Behrens, H. M. et al. Pyocin S5 Import into Pseudomonas aeruginosa Reveals a Generic Mode of Bacteriocin Transport. MBio 11, (2020).

17. Elfarash, A. et al. Pore-forming pyocin S5 utilizes the FptA ferripyochelin receptor to kill Pseudomonas aeruginosa. Microbiology 160, 261–269 (2014).

18. Elfarash, A., Wei, Q. & Cornelis, P. The soluble pyocins S2 and S4 from Pseudomonas aeruginosa bind to the same FpvAI receptor. Microbiologyopen 1, 268–75 (2012).

19. Denayer, S., Matthijs, S. & Cornelis, P. Pyocin S2 (Sa) kills Pseudomonas aeruginosa strains via the FpvA type I ferripyoverdine receptor. J. Bacteriol. 189, 7663–8 (2007).

20. McCaughey, L. C. et al. Discovery, characterization and in vivo activity of pyocin SD2, a protein antibiotic from Pseudomonas aeruginosa. Biochem. J. 473, 2345–2358 (2016).

21. Baysse, C. et al. Uptake of pyocin S3 occurs through the outer membrane ferripyoverdine type II receptor of Pseudomonas aeruginosa. J. Bacteriol. 181, 3849–3851 (1999).

22. Michel-Briand, Y. & Baysse, C. The pyocins of Pseudomonas aeruginosa. Biochimie vol. 84 499–510 (2002).

23. McCaughey, L. C., Ritchie, N. D., Douce, G. R., Evans, T. J. & Walker, D. Efficacy of species-specific protein antibiotics in a murine model of acute Pseudomonas aeruginosa lung infection. Sci. Rep. 6, 30201 (2016).

24. Chousterman, B. G., Swirski, F. K. & Weber, G. F. Cytokine storm and sepsis disease pathogenesis. Seminars in Immunopathology vol. 39 517–528 (2017).

25. Singer, M. et al. The third international consensus definitions for sepsis and septic shock (sepsis-3). JAMA - Journal of the American Medical Association vol. 315 801–810 (2016).

